# Mathematical investigation of microbial quorum sensing under various flow conditions

**DOI:** 10.1101/2020.01.09.900027

**Authors:** Heewon Jung, Christof Meile

## Abstract

Microorganisms efficiently coordinate phenotype expressions through a decision-making process known as quorum sensing (QS). We investigated QS amongst heterogeneously distributed microbial aggregates under various flow conditions using a process-driven numerical model. Model simulations assess the conditions suitable for QS induction and quantify the importance of advective transport of signaling molecules. In addition, advection dilutes signaling molecules so that faster flow conditions require higher microbial densities, faster signal production rates, or higher sensitivities to signaling molecules to induce QS. However, autoinduction of signal production can substantially increase the transport distance of signaling molecules in both upstream and downstream directions. We present approximate analytical solutions of the advection-diffusion-reaction equation that describe the concentration profiles of signaling molecules for a wide range of flow and reaction rates. These empirical relationships, which predict the distribution of dissolved solutes following zero-order production kinetics along pore channels, allow to quantitatively estimate the effective communication distances amongst multiple microbial aggregates without further numerical simulations.

**Author Summary:** Microbes can interact with their surrounding environments by producing and sensing small signaling molecules. When the microbes experience a high enough concentration of the signaling molecules, they express certain phenotypes which is often energetically expensive. This microbial decision-making process known as quorum sensing (QS) has been understood to confer evolutionary benefits. However, it is still not completely understood how transport of the produced signaling molecules affects QS. Using a mathematical approach investigating QS across a range of environmentally relevant flow conditions, we find that advective transport promotes QS downstream yet also dilutes the concentration of signaling molecules. We quantify the importance of microbial cell location with respect to both other microbes and flow direction. By analyzing complex numerical simulation results, we provide analytical approximations to assess the distribution of signaling molecules in pore channels across a range of flow and reaction conditions.

## Introduction

Microorganisms preferentially reside on solid surfaces, which often leads to a closer proximity of neighboring cells than when in a planktonic form [1]. At elevated cell densities, microorganisms need to efficiently coordinate the expression of energetically expensive phenotypes, such as biofilm development, exoenzyme production, and microbial dispersal. Efficiency is achieved by producing and detecting relatively cheap signaling molecules which regulate the phenotype expression only when a sufficient signal concentration has been reached [2]. This microbial decision-making process called “quorum sensing (QS)” was originally understood as a cell-to-cell communication to identify conspecific population density and accomplish cooperative behaviors [3]. However, a number of studies have indicated that QS is not necessarily a social trait [4,5] and depends not only on the population but also on the spatial distribution of microbial cells [6,7]. These observations led to an alternative QS concept in which QS depends strictly on the local concentration of signaling molecules [8,9]. This suggests that, to understand QS processes, an integrative approach is required analyzing a multitude of factors including microbial density [3], production and decay kinetics [10,11], and transport of signaling molecules through advection and diffusion [4], as well as the spatial distribution of microorganisms [7]. Thus, spatial constraints and responses may be as important as other biological considerations for the evolution and maintenance of QS. This idea is known to be true in biofilms where cooperative strategies are able to evolve if cooperators are spatially aggregated [12].

Individual microbial cells synthesize and release signaling molecules at a basal rate. At low population densities, the concentration of signaling molecules remains low as it degrades both biotically and abiotically [11,13]. At a sufficiently high microbial population density, however, the extracellular concentration of signaling molecules reaches a threshold concentration that activates gene and phenotypes expression [9]. When QS regulates the production of costly public goods, this balances production cost and the overall benefit [27,35,36], while under nutrient limited conditions, QS can regulate microbial dispersal (e.g. [37,38]), improving chances of survival. QS induction also often upregulates genes controlling production of signaling molecules resulting in enhanced signal production [10,14]. Such autoinduction has been thought to confer evolutionary stability and fitness advantages [15–17], but its effects on neighboring microbial aggregates and evolutionary benefits in a spatial context have not be fully understood.

QS induction is affected by mass transport characteristics controlling the spatial distribution of signaling molecules. In a confined space, even a single microbial cell can be QS induced if the signaling molecules accumulate to sufficiently high concentration in the confined space [5]. However, higher population densities are required for QS induction in a large open space because the signaling molecules are diluted due to diffusive loss to the surrounding medium [7,18]. Advection may dilute the signaling molecules more effectively than diffusion and repress QS induction [19–23]. The signaling molecules transported either via advection or diffusion can induce QS in neighboring cells if they receive enough signaling molecules. The distance between two QS induced microbial cells or aggregates is referred to as the “calling distance” [24], and has been reported to be 5 - 78 μm between individual cells [24] and ∼180 μm between microbial aggregates [25]. The dependence of QS processes on advection and diffusion suggests that transport regimes affect calling distances, highlighting the importance of relative positioning of microorganisms coupled with the mass transport characteristics of a habitat.

Here, we evaluate the effect of combined diffusive and advective transport on QS processes in environmentally relevant conditions using a reactive transport modeling approach. The advection-diffusion-reaction equation (ADRE) was nondimensionalized to capture the characteristic properties of QS systems (i.e. effects of the rate of signaling molecule production, mass transport, and spatial distribution of microbial aggregates) and used to formulate empirical expressions describing concentration profiles of signaling molecules under various flow conditions. Using these relationships, we evaluate calling distances and threshold biochemical conditions for QS induction of a single microbial aggregate under various flow conditions. Then, we investigate QS interactions between heterogeneously distributed microbial aggregates. Finally, we demonstrate the importance of autoinduction for coordinated microbial behaviors in advection-dominated environments.

## Process description

The distribution of signaling molecules is described by

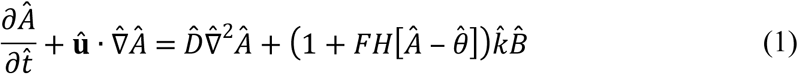

where 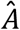 is a concentration of signaling molecules, **û** is the flow velocity vector representing parabolic Poiseuille flow (see the section “Numerical method”), and 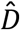 is the molecular diffusivity. The last term on the right-hand side of the equation represents the production rate of signaling molecules. QS induction often displays a switch-like behavior [9,26,27], which is represented in the model by a step function with a higher signal production rate above the threshold concentration of signaling molecule:

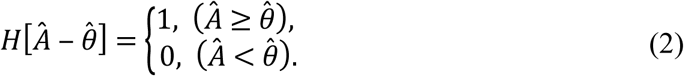

and *F* represents a multiplication factor. Here *F* is set to either 10 to reflects the magnitude of autoinduced signal production [10] or 0 in the absence of autoinduction. 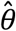 is the QS induction threshold, 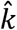 is the basal production rate constant of signaling molecules, and 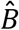 is the microbial density. Note that we are ignoring the breakdown of signaling molecules [28], limiting us to settings where production and transport are the dominant processes.

To describe the characteristic properties of a microbial system across various flow and reaction conditions, Eq. 1 was recast by introducing dimensionless quantities 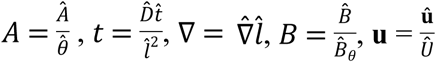, where 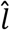 is a characteristic length (i.e. the width of the flow channel), *Û* is a characteristic fluid velocity (here, the average pore fluid velocity), and 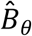 is a threshold biomass density required for QS induction, resulting in:

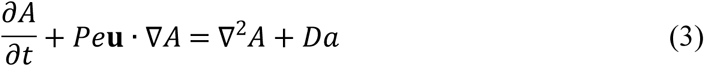

This nondimensionalized ADRE is fully characterized by the Péclet number, expressing the magnitude of advective flow relative to diffusion 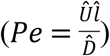, and the diffusive Damköhler number, comparing reaction to diffusion (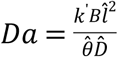; where 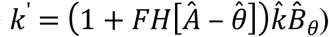. A system with high *Da* - either due to high *k*’ (i.e. fast signal production), high *B* (i.e. high microbial density), or low 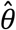 (i.e. high sensitivity to signaling molecules) - is more likely to be QS induced.

An important property of Eq. 3 is that its solution linearly scales in *Da*. For example, if *Da* is increased 2-fold at a fixed *Pe* condition, the concentrations of signaling molecule are doubled. This linearity allows to calculate the concentration distribution of signaling molecules for any *Da* from a single simulation result with an arbitrary *Da* at a given *Pe*. However, this simple approach cannot be applied to the flow conditions because the solution is not linear in *Pe*. Therefore, multiple numerical simulations were carried out with 24 *Pe* conditions (*Pe* ∈ {0.5, 0.6, 0.7, 0.8, 0.9, 1, 1.5, 2, 2.5, 3, 3.5, 4, 4.5, 5, 5.5, 6, 6.5, 7, 7.5, 8, 8.5, 9, 9.5, 10}) while *Da* was fixed at 5. For the 2D simulations in a pore channel shown below, the flow field was established by imposing pressures at in- and outlet and no flow conditions at the top and bottom boundaries, resulting in a flow from left to right. Fixed concentration (*A*|_left-boundary, x=0_ = 0) and no-gradient (*∂A*/*∂x*|_right-boundary, x=4_ = 0) boundary conditions were imposed at the inlet and outlet boundaries, with no-flux at the top and bottom boundaries, respectively. All simulations were run to steady state.

## Results and discussion

### QS processes of a single microbial aggregate

The effect of various flow conditions on the distribution of signaling molecules (*A*) produced from a single microbial aggregate located at *x* = 1 was investigated under various *Pe* conditions (0.5 ≤ *Pe* ≤ 10) while *Da* was fixed at 5 (Fig. 1). The QS induction enhancing the signal production rate was not considered.

**Fig. 1.**
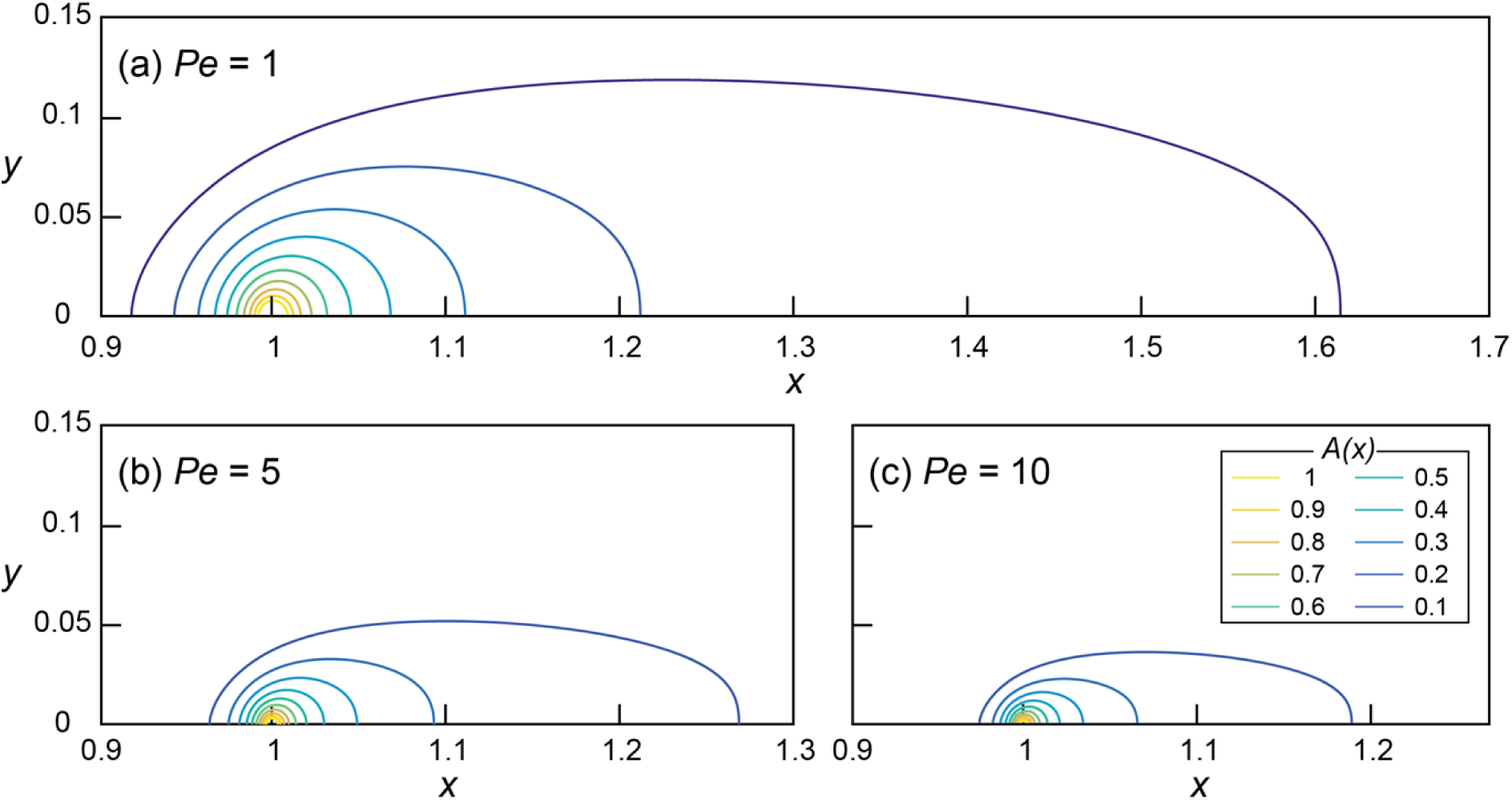
Concentration fields of signal concentration *A* at *Da* = 5 and (a) *Pe* = 1, (b) *Pe* = 5, and (c) *Pe* = 10, without autoinduction (*F* = 0). Note the difference in scale on the horizontal axis.

The signal concentration fields developed under various advective flows show maximum concentrations (*A*_*max*_ = *A*(*x*=1)) decreasing with increasing *Pe* (i.e. faster advective flow): *A*_*max*_ decreased from 1.68 (*Pe* = 1) to 1.35 (*Pe* = 5) and 1.21 (*Pe* = 10). However, *A*_*max*_ of all of the simulations with *Da* = 5 exceeded 1(i.e. 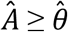), indicating the potential for QS induction. The threshold *Da* for QS induction (*Da*_θ_), where *A*_*max*_ = 1, can easily be computed using the linearity of the nondimensionalized ADRE in *Da* (Eq. 3). For example, *Da*_θ_ at *Pe* = 1 was calculated by dividing *Da* = 5 by its corresponding *A*_*max*_ = 1.68 which resulted in *Da*_θ_ = 2.98. Thus, at *Pe* = 1, conditions for which *Da* ≥ 2.98 lead to or exceed the concentration of signaling molecules needed for QS induction. Fig. 2 shows the calculated *Da*_θ_ for each simulated *Pe* condition.

**Fig. 2.**
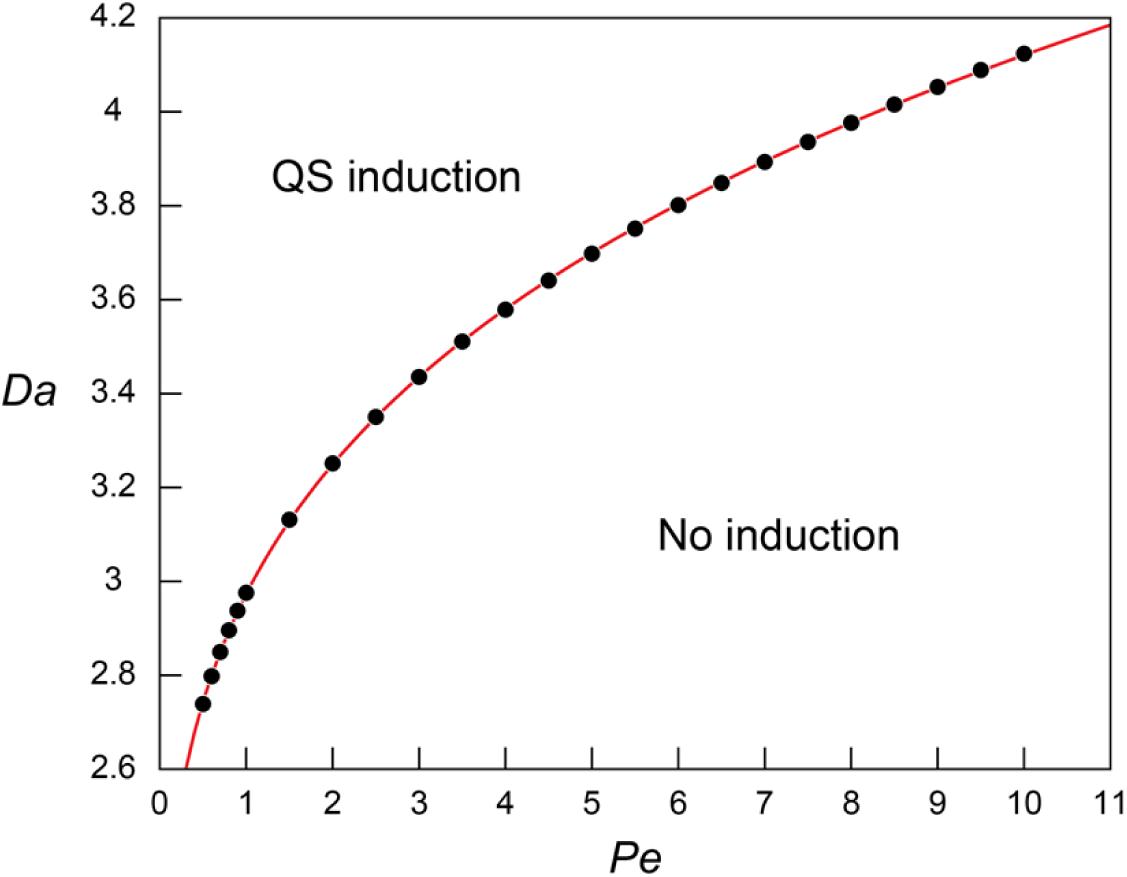
The relationship between the threshold *Da* for QS induction (*Da*_θ_) and *Pe*. The simulation results (block dots) were fitted using the power regression (red line; Eq. 4).

The regression analysis revealed that the simulated *Da*_θ_ for QS induction varies as a function of *Pe* following the power law:

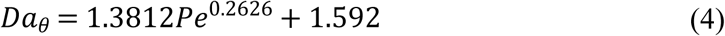

The increasing *Da*_θ_ along with the increasing *Pe* indicates higher *Da* (i.e. higher microbial density (*B*), higher signal production rate constant (*k*’), or lower QS induction threshold 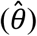) is required for QS induction under higher *Pe*. This result corresponds to the observed repressed QS induction under the presence of advection [19,21,23]. Eq. 4 was further evaluated by applying the experimentally measured QS parameters of *Pseudomonas putida* (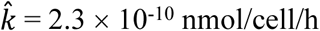, and 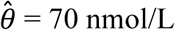 [10]) in a flow system where 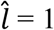 cm and 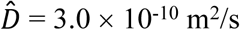 [29]. Our results show 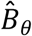 of 9.77, 12.2, and 13.5 × 10^6^ cells/mL at *Pe* = 1, 5, and 10, respectively. If Eq. 4 is extrapolated to diffusion only transport condition (*Pe* = 0, *Da*_θ_ = 1.592), 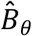 is estimated as 5.23 × 10^6^ cells/mL which largely agree with the experimental observation of 2.69 ∼ 6.23 × 10^6^ cells/mL where signal concentration starts to show a strong spike (Table S1 in [10]).

In addition to reducing *A*_*max*_, advection also influenced the spatial distribution of signaling molecules. We define the “transport distance” (*d*) as the distance between the point of production (*x*_0_) and the point (*x*_1_) where the signal concentration reaches a certain value *A*\* (i.e. *d* = |*x*_0_ – *x*_1_|), distinguishing it from the “calling distance” between two QS induced microbial cells or aggregates. If the signal transport occurred only through diffusion, transport distances would be isotropic [7]. However, advection resulted in anisotropic concentration distribution where upstream transport distances (*d*_*up*_) are much shorter than the downstream distances (*d*_*dn*_). Moreover, fast advective flows (i.e. high *Pe*) reduced overall transport distances which are illustrated in Fig. 1a-c as the shrinking areas covered by contour lines. For example, the (nondimensional) transport distances to the location where *A*_1_ = 0.4 are *d*_*up*_ = 0.035 and *d*_*dn*_ = 0.068 at *Pe* = 1 and *Da* = 5 (Fig. 1a). These values decrease to *d*_*up*_ = 0.018 and *d*_*dn*_ = 0.043 at *Pe* = 5 (Fig. 1b) and to *d*_*up*_ = 0.013 and *d*_*dn*_ = 0.036 at *Pe* = 10 (Fig. 1c).

### Empirical approximation of concentration profiles

Obtaining transport distances for different *Pe* conditions requires running numerical simulations for each of the corresponding *Pe*. However, this may be avoided if we can express the concentration profiles as a function of *Pe*. For this purpose, parametric regression analysis was applied to the numerically obtained concentration profiles along the bottom of the flow channel (Fig. 3).

**Fig. 3.**
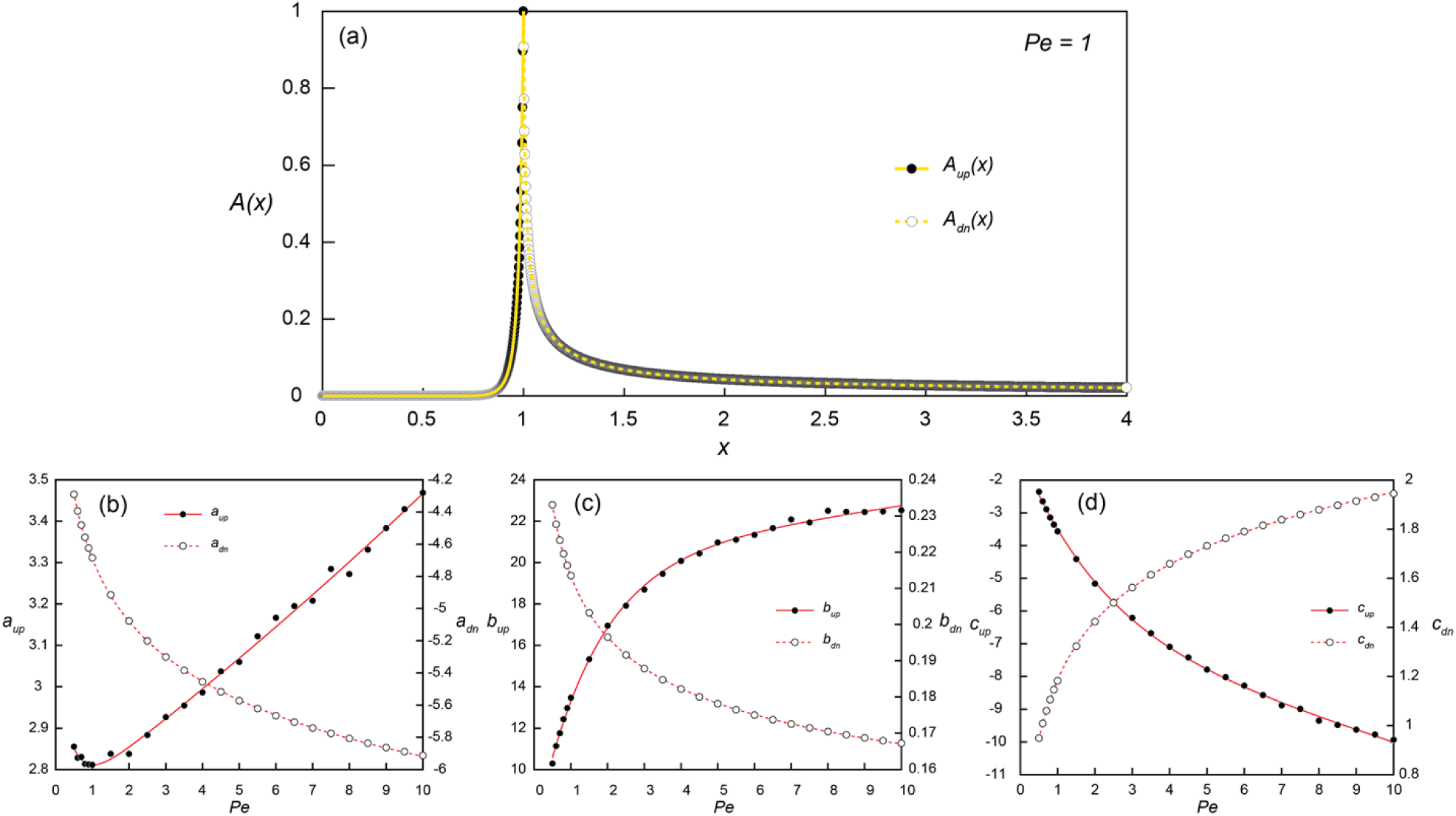
(a) Simulated (dots) and reconstructed (lines) concentration profiles along the bottom of the flow channel (*y* = 0.001) at *Pe* = 1 and *Da* = *Da*_θ_ = 2.98. The upstream (*x* ≤ 1; *A*_*up*_(*x*); solid line) and downstream (*x* > 1; *A*_*dn*_(*x*); dashed line) concentration profiles were obtained from Eqs. 5 and 6, respectively. (b-d) The coefficients for *A*_*up*_(*x*) (*a*_*up*_, *b*_*up*_, and *c*_*up*_) and *A*_*dn*_(*x*) (*a*_*dn*_, *b*_*dn*_, and *c*_*dn*_) obtained from the parametric regressions of the simulated concentration profiles at each simulated *Pe* conditions with Eq. 5 (black dots) and Eq. 6 (white dots), respectively. The solid and dashed lines are the exponential (Eqs. 7-9) and power fits (Eqs. 10-12) of the estimated coefficients as a function of *Pe*.

Several parametric regression models (linear, power, exponential, and polynomial models) were tested to the upstream (*A*_*up*_(*x*); 0 ≤ *x* ≤ 1) and downstream (*A*_*dn*_(*x*); 1 < *x* ≤ 4) signal concentration profiles. Among the tested regression models, the exponential (Eq. 5) and power-law models (Eq. 6) provided the best fit for log-transformed upstream and downstream signal concentration profiles, respectively. In the regression analysis of upstream profiles, only the locations where *A(x)* > 0.001 were used to improve the fitting quality and the signal concentration at *x* = 1 was fixed as 1. The additional regression analysis was then carried out for the coefficients (*a, b*, and *c*) obtained from simulated profiles at 24 *Pe* conditions to construct a relationship between the coefficients and *Pe* (Fig. 3b-d). The exponential and power-law models provided the best fit for the upstream (Eqs. 7-9) and downstream coefficients (Eqs. 10-12), respectively:

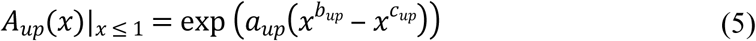

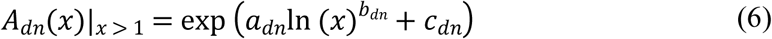

where *A*_*up*_ and *A*_*dn*_ are 0 in the down- and up-stream directions, respectively, and

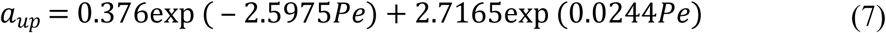

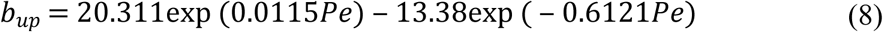

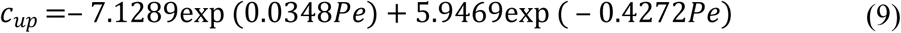

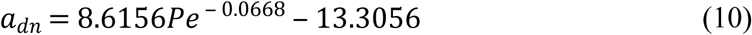

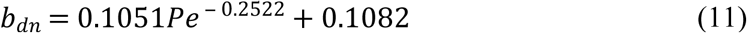

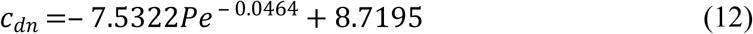

Eqs. 5 and 6 can be used as approximations of the concentration profiles along a pore channel without running simulations for various *Pe* conditions, with the microbial aggregate located at *x* = 1. Due to the linearity in *Da*, the concentration profiles at different *Da* conditions can be calculated simply by multiplying *Da*/*Da*_θ_ to Eqs. 5 and 6, so that

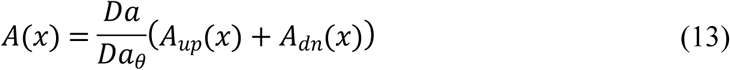

These analytical expressions are applicable not only to QS but also to other chemical processes subject to zero-order production reactions. They are valid when the microbial aggregates have a negligible impact on flow fields, which is a reasonable approximation for low microbial density conditions. However, it may not hold when biomass grow large and perturb flows substantially which would require models fully resolving nonlinear feedback between fluid flows and transport phenomena [30–32]. The equations become less accurate at low *Pe* as under low flow conditions, the estimates from Eq. 13 in a flow channel with a small width (i.e. low 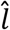 and *Pe*) could underestimate the actual concentration because the confined channel width would push the produced chemical further upstream and downstream.

### The effect of QS induced signal production on transport distances

QS often involves autoinduction which substantially increases signal production rates. The effect of autoinduction on transport distances was investigated by using Eq. 13 for the conditions without (*F* = 0; *Da* = *Da*_θ_) and with (*F* = 10; *Da* = 11*Da*_θ_) enhanced signal production. The transport distances from a single microbial aggregate under various *Pe* were then calculated using Eq. 13 for the location *x*.

Fig. 4 shows the transport distances without (Fig. 4a) and with (Fig. 4b) the enhanced signal production at *Pe* = 1, 5, and 10. The concentration ratios (0.1 ≤ *A*/*A*_*max*_ ≤ 0.9) were used instead of absolute concentrations to generalize transport distances for various *Da* conditions. For example, the transport distance (*d*_*A*_) for *A*/*A*_*max*_ = 0.5 indicates that *A*(*x*_0_+*d*_*A*_) = 0.5 if *Da* = *Da*_θ_ while *A*(*x*_0_+*d*_*A*_) = 0.05 when *Da* = 0.1*Da*_θ_. The consequence of the enhanced signal production was the significant increase of *d*_*up*_ and *d*_*dn*_. Without the enhanced signal production, *d*_*up*_ and *d*_*dn*_ for *A/A*_*max*_ = 0.4 at *Pe* = 1 were estimated as 0.021 and 0.024, respectively (Fig. 4a). These values increased to *d*_*up*_ = 0.1 and *d*_*dn*_ = 1.28 with the enhanced signal production (Fig. 4b). The downstream transport distance of 1.28 is translated into 6.4 mm in a flow channel with 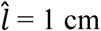. This result is much longer than the generally observed ranges of calling distances [33]. However, we emphasize again that the transport distance merely indicates the distance of signaling molecules transported from a source location while the calling distance involves QS induced microbial cells or aggregates.

**Fig. 4.**
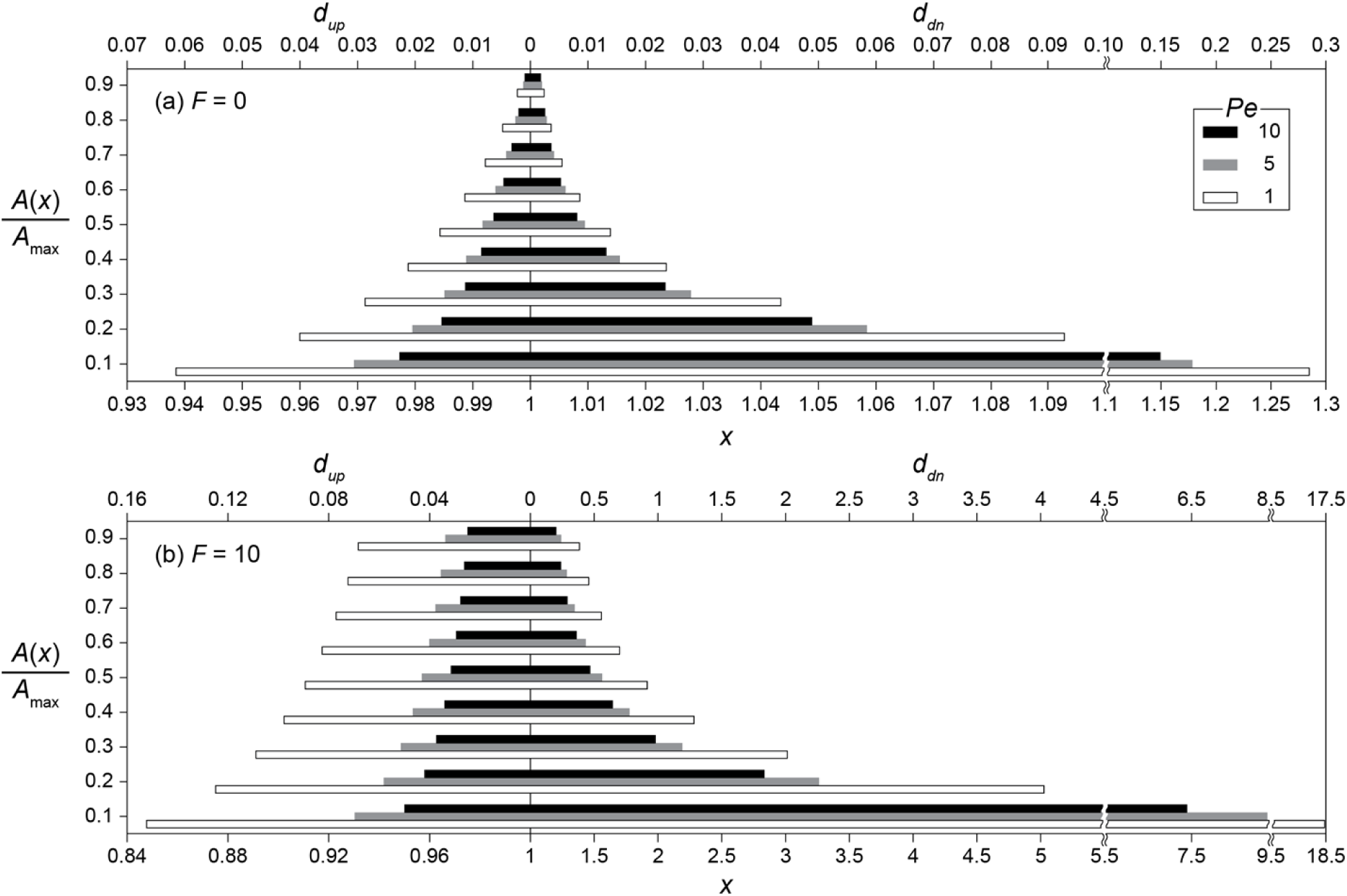
Upstream (*d*_*up*_) and downstream (*d*_*dn*_) transport distances (a) without (*F* = 0) and (b) with (*F* =10) enhanced signal production for the concentration ratios (0.1 ≤ *A*/*A*_*max*_ ≤ 0.9) at *Pe* = 1, 5, and 10. Note the different scale between panels (a) and (b), and between the distances up- and downstream from 0 (*d*_*up*_ and *d*_*dn*_) in (b).

### QS induction between spatially distributed multiple microbial aggregates

QS processes of multiple aggregates were investigated by constructing the concentration profiles using Eq. 14. Concentration fields of signaling molecules with multiple microbial aggregates can be calculated as the superposition of the concentration profile produced by each individual aggregate:

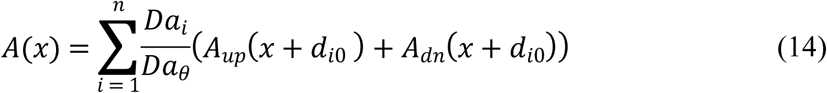

where *n* is the number of aggregates, *d*_*i0*_ is the distance between *x*_*i*_ and *x*_*0*_ (*d*_*i0*_ = *x*_*i*_ – *x*_*0*_), *x*_*i*_ is the location of *i*th aggregate, *x*_*0*_ is the reference location (*x*_*0*_ = 1), *Da*_i_ is the *Da* calculated only with the density of *i*th microbial aggregate (i.e. microscopic *Da*), and *A*_*up*_ and *A*_*dn*_ are Eqs. 5 and 6, respectively. Here, an example system with macroscopic *Da (Da*_T_ = Σ*Da*_i_) = 3.2*Da*_θ_ consist of four aggregates (𝔸_1-4_) located at *x*_1_ = 0.4, *x*_2_ = 1, *x*_3_ = 1.096, and *x*_4_ = 1.7 with the evenly distributed microscopic *Da*_i_ (i.e. *Da*_1_ = *Da*_2_ = *Da*_3_ = *Da*_4_ = 0.8*Da*_θ_) was tested. In using Eq. 14, the profile was first constructed for *Da*_*i*_ *= Da\** that does not consider autoinduction (*F* = 0). Then, if there is an aggregate with *A*(*x*_i_) ≥ 1, the profile was recalculated with updated *Da*_i_ = (1+*F*)×*Da*\* until all *Da*_i_ with *A* ≥ 1 were updated.

The signal concentration profile produced by four aggregates without the enhanced signal production (*F* = 0) illustrates the crucial importance of relative positioning of microbial aggregates for QS induction with respect not only to each aggregate but also to the flow direction (Fig. 5a). The microscopic *Da*_i_ was set such that the maximum concentration produced by a single aggregate was 0.8, as observed at the most upstream location (𝔸_1_ at *x*_*1*_ = 0.4). But due to transport, the local concentration at 𝔸_2_ reached 0.879, receiving *A* of 0.048 and 0.031 from 𝔸_1_ and 𝔸_3_, respectively. 𝔸_3_ received slightly less signaling molecules from 𝔸_1_ (*A* = 0.044) due to the longer distance of 𝔸_3_ than 𝔸_2_ from 𝔸_1_. However, 𝔸_2_ provided much more signaling molecules (*A* = 0.157) to 𝔸_3_ than was provided by 𝔸_3_ because of advective flows favoring downstream transport of signaling molecules (Figs. 2 and 4). As a consequence, 𝔸_3_ exceeded the QS threshold (*A*(*x*_3_) = 0.044 from 𝔸_1_ + 0.157 from 𝔸_2_ + 0.8 from 𝔸_3_ + 0 from 𝔸_4_ = 1.001 > 1) while the upstream located 𝔸_2_ did not. The QS induction of 𝔸_3_ demonstrates the importance of transport distances. QS induction was achieved because of the upstream aggregates located within the transport distance of 0.696. However, the calling distance would have been estimated as the length of a grid voxel (0.002) because only 𝔸_3_ was QS induced. Therefore, considering only the calling distance could lead to the wrong conclusion that the local *Da* condition at 𝔸_3_ (i.e. *Da*_3_ = 0.8*Da*_θ_) is a sufficient condition for QS induction. Although 𝔸_4_ did not reach the QS induction threshold, it received *A* from all the other aggregates resulting in a concentration (*A*(*x*_4_) = 0.029 + 0.044 + 0.048 + 0.8 = 0.921) that was higher than at 𝔸_2_ despite the longest separation distance from other aggregates.

**Fig. 5.**
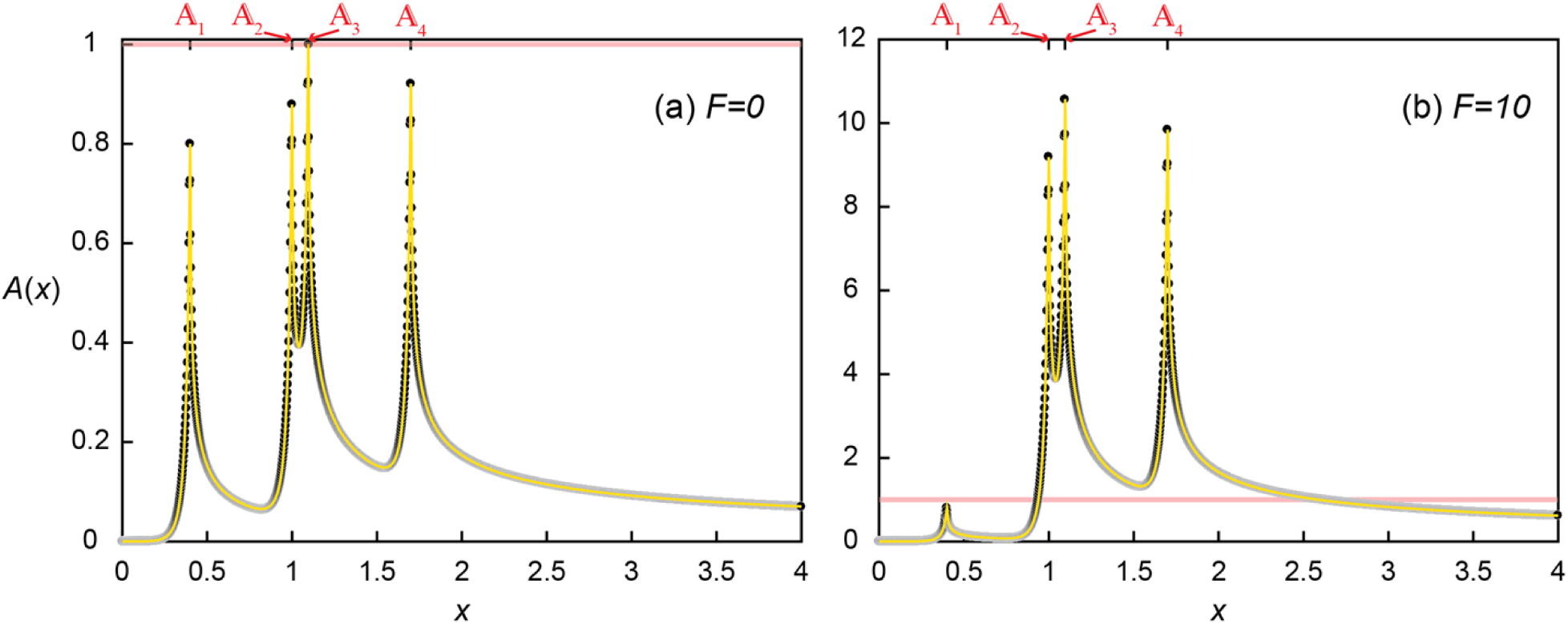
Concentration profile (a) without (*F* = 0) and (b) with (*F* = 10) enhanced signal production where four aggregates are located at *x*_1_ = 0.4, *x*_2_ = 1, *x*_3_ = 1.096, and *x*_4_ = 1.7. Black dots are the simulated results and the yellow lines represent the profile from Eq. 14.

Accounting for QS induction (*F* = 10) increased the transport distances and hence induced other aggregates (Fig. 5b). With the same spatial distribution, QS-induced 𝔸_3_ produced signaling molecules much more and faster (i.e. 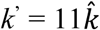 and *Da*_3_ = 8.8*Da*_θ_) and provided more signaling molecules to 𝔸_2_. As a result, *A*(*x*_2_) exceeded the QS threshold (0.048 + 0.8 + 0.335 + 0 = 1.183). The QS induction of 𝔸_2_ and 𝔸_3_ resulted in the final signal concentrations of *A*(*x*_2_) = 9.183 (= 0.048 + 8.8 + 0.035 + 0) and *A*(*x*_3_) = 10.569 (= 0.044 + 1.725 + 8.8 + 0). While 𝔸_4_ still did not contribute signaling molecules to any of upstream aggregates, enhanced contribution from 𝔸_2_ and 𝔸_3_ QS induced 𝔸_4_, *A*(*x*_4_) = 9.839 (= 0.029 + 0.48 + 0.53 + 8.8). Despite increased transport distances by QS induction, 𝔸_1_ was still too far away from the other aggregates thus the signal concentration at 𝔸_1_ remained unchanged *A*(*x*_1_) = 0.8. As a result of the QS induction of 𝔸_2-4_, *Da*_T_ had increased from the initial 3.2*Da*_θ_ (= 0.8*Da*_θ_ × 4) to 27.2*Da*_θ_ (= 0.8*Da*_θ_ + 3 × 11 × 0.8*Da*_θ_).

This example illustrates the importance of enhanced signal production on the spatial propagation of QS induction. While only 𝔸_3_ experienced signaling molecule levels that could induce QS when all the aggregates produce signaling molecules at the basal production rate, the enhanced signal production of 𝔸_3_ when considering induced production (*F* = 10) provided more signaling molecules to its adjacent microbial aggregates and resulted in the QS induction of neighboring aggregates, 𝔸_2_ and 𝔸_4_. It may be counterintuitive that the upstream-located 𝔸_2_ was also QS-induced by the contribution from 𝔸_3_ despite the contracted upstream transport distances under the presence of advective flows. This result shows that the enhanced signal production can overcome the influence of advection and promote QS induction, and provide a way to provoke upstream microbial aggregates, for example, to slow down the substrate consumption to ensure efficient resource utilization in crowded environments [34].

### Implications

This study has demonstrated that advection and the enhanced signal production can determine the spatial extent of QS induction. Although QS mediated gene expression has been understood as evolutionarily beneficial collective behaviors, long transport distances observed in study suggests that it may not be always true. The transport of signaling molecules, especially in downstream direction, combined with enhanced signal production, suggests that QS induction can be decoupled from microbial density. In the above example, any microbial cell located where *A* > 1 (e.g. *A*(*x* = 2.5) = 1.05 in Fig. 5b) would have been QS-induced, independent of the local cell density. This could lead to detrimental impacts on a microbial population, unless there are other counteracting mechanisms such as differential QS induction sensitivity to signal concentration even in within a clonal population [25] or biofilm formation modifying local transport characteristics [39]. Future investigations should explicitly examine the evolutionary consequences of QS strategies in spatially heterogeneous environments under advective-diffusion-reaction dynamics.

### Numerical methods

We used the Lattice Boltzmann (LB) method to implement a numerical model for the transport of signaling molecules due to diffusion and advection. The LB method is a mesoscopic approach solving the Boltzmann equation across a defined set of particles which recovers the macroscopic Navier-Stokes equation (NSE) and ADRE [40]. First, we obtained the flow field by solving the particle distribution function *f*:

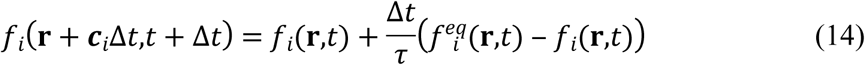

where particles *f*_*i*_ (**r**,t) travel in the direction *i* with the lattice velocity ***c***_*i*_ (***c***_*0*_ = (0,0), ***c***_*1*_ = (1,0), ***c***_*2*_ = (0,1), ***c***_*3*_ = (−1,0), ***c***_*4*_ = (0,-1), ***c***_*5*_ = (1,1), ***c***_*6*_ = (−1,1), ***c***_*7*_ = (−1,-1), ***c***_*8*_ = (1,-1)) to a new position **r**+ ***c***_*i*_Δt after a time step Δt. The relaxation time (τ) was described by the commonly used Bhatnagar-Gross-Krook collision operator [41] and the D2Q9 lattice with the corresponding equilibrium distribution function:

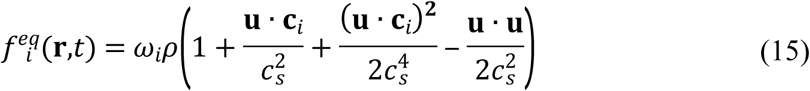

where *ω*_*i*_ are lattice weights (*ω*_*0*_ = 4/9, *ω*_*1-4*_ = 1/9, *ω*_*5-8*_ = 1/36), *c*_*s*_ is a lattice dependent constant (here, 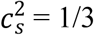), and **u** is the macroscopic flow velocity. The moments of the discretized mesoscopic particles retrieve the macroscopic density (ρ = Σ *f*_*i*_) and momentum (ρ**u** = Σ ***c***_*i*_ *f*_*i*_). The Chapman-Enskog expansion showed that this LB approach recovers the incompressible NSE with the viscosity 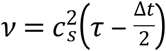 [40].

Once the flow field was obtained, we simulated solute transport with a particle distribution function *g*, using the regularized LB algorithm (RLB) for numerical accuracy [42,43] and the D2Q5 lattice for numerical efficiency [44]:

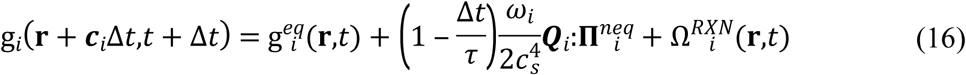

where ***c***_*i*_ are the lattice velocities (***c***_*0*_ = (0,0), ***c***_*1*_ = (1,0), ***c***_*2*_ = (0,1), ***c***_*3*_ = (−1,0), ***c***_*4*_ = (0,-1), ***c***_*5*_ = (1,1)) corresponding to the lattice weights *ω*_*i*_ (*ω*_*0*_ = 1/3, *ω*_*1-4*_ = 1/6), and 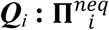 is the tensor contraction between the two tensors 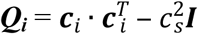 and 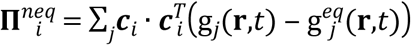.

The reaction term in the Eq. 3 describes the production of signaling molecules:

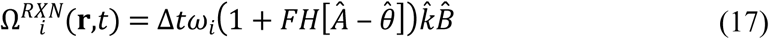

With the imposed flow field from Eq. 1, the LB transport solver (Eq. 3) recovers the ADRE (Eq. 1) with the diffusivity 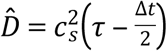.

Simulations were conducted for a 2D flow channel of non-dimensional length of 4 and a width of 2, discretized with 2000 × 1000 grid elements. The flow field (Eq. 1) was generated by imposing fixed pressures at inlet (*x* = 0) and outlet (*x* = 4) with no flow boundaries in both normal and tangential direction at the bottom (*y* = 0) and top (*y* = 2) of the domain resulting in parabolic Poiseuille flows. Simulations were carried out under low Mach numbers (Ma ≪ 1) to ensure incompressible flow conditions [40].

## Acknowledgement

We thank Chris Kempes for comments on the manuscript. The LB code of this study is available at https://bitbucket.org/MeileLab/jung_qsTpDistn.

## Author Contributions

Conceived and designed the experiments: HJ CM. Performed the experiments: HJ. Analyzed the data: HJ. Contributed reagents/materials/analysis tools: HJ CM. Wrote the paper: HJ CM.

